# Insights on Poster Preparation Practices in Life Sciences

**DOI:** 10.1101/2020.10.08.331413

**Authors:** Helena Klara Jambor

## Abstract

Posters are intended to spark scientific dialogue and omnipresent at biological conferences. Guides and how-to articles help life scientists in preparing informative visualizations in poster format. However, those posters shown at conferences are at present often overloaded with data and text and are poorly designed, which in sum hinder, rather than spark communication. Here, we surveyed life scientists themselves to understand why posters are such a struggle and to shed light on poster preparation from their subjective perspectives. We find that biologist spend on average two entire days preparing one poster, with half of the time devoted to visual design aspects. Most respond that they receive no design or software training and also receive little to no feedback when preparing their visualizations. In conclusion, our data suggests that some training, beyond providing literature, of life scientists in visualization principles and tools would likely improve conference poster quality. In addition, this would also improve other common visuals such as figures and slides, and thus overall benefit the science communication of researchers.

## Introduction

Visualizations are important to communicate biological and medical sciences since antiquity (Stückelberger, 1994). Today, scientists visually present biomedical data in manuscript figures, talk slides and, at least since the 1970ies, demonstrably also with scientific posters at conferences (Dubois, 1985a; Maugh, 1974). Poster presentation sessions are a popular event (Rowe, 2017), especially so when participant numbers exceed the speaker slots. At large meetings thousands of posters may be presented: 2500 at the meeting of the American Society for Cell Biology, 3000 at the meeting of the American Society for Human Genetics, and 1500 at the meeting of Technology Association of Grantmakers (TAG). Posters are particularly popular among early career researchers, who use posters to show in-progress projects and have personal discussions. For many early career scientists, a poster session today is their first experience in publicly presenting results to colleagues and a wider audience.

Posters are, similar to infographics, a large-scale visualization format juxtaposed of explanatory text, pictures, schematics, and/or data visualizations, charts, or diagrams. Across all of life sciences, posters include a header with title and authors information, an overview of the rationale, methods, and results, and at times references, acknowledgements, and summaries (Brown, 1996; Dubois, 1985b). This organization was previously described as the “IMRAD format”, and echoes the organisation of a scientific manuscript, with a title including the authors followed by introduction, oftentimes the text submitted as abstract, a brief methodology section, representative results, and a brief discussion (Brown, 1996). Just like in an infographic, the reading orientation dictates in which order the elements of the posters should be read and must be well designed by the author (Dubois, 1985b). At poster sessions, scientists should provide audiences a poster presentation, a brief narration of the entire poster also referred to as the elevator pitch (Erren and Bourne, 2007). The poster, along with its presentation, thus is a form of visually-aided storytelling.

Several guidelines and how-to papers have been published that aim to help life science researchers in designing a legible and concise poster (Barker and Phillips, 2021; Block, 1996; Brown, 1997, 1996; Erren and Bourne, 2007; Wang et al., 2022). These guidelines are explicitly written for biologists with little or no previous knowledge in visual design and low visualization literacy. Boullata et al. and also Gundogan et al provide updates with an exemplar template, details on text, font size, information flow and the importance of feedback (Boullata and Mancuso, 2007; Gundogan et al., 2016). Erren and Bourne, 2007, expand from the display and also provide recommendations for the actual oral presentation and suggest to having an “elevator pitch” ready. In addition, there is also a growing body of literature helping life scientists with the overall presentation of visual information. Bang Wong initiated a long-running series of articles that cover many aspects of design and visualization principles that are relevant also to posters or to the elements within a poster: these articles comment on the elements of visual style, layout and Gestalt principles, but also colours and storytelling (Krzywinski, 2013; Krzywinski and Cairo, 2013; Wong, 2011a, 2011b, 2011c). Importantly, the intended audiences are early career life scientists struggling with visual design of information. Helping biologists further, many universities already provide poster templates for standard software that have a pre-set layout and new tools such as Biorender have ready-to-go poster templates.

Despite these guidelines, not all posters are well-designed to be easily understandable. A commentary recount of all that can go wrong with poster presentations is given by Thomas Wolcott (Wolcott, 1997). He lists that posters are not self-explanatory and hard to read, have too much text, cryptic abbreviations, inconsistent colours and a lack of visual order. A study recently investigated this quantitatively. The authors reveal that the format of award-winning posters at life science conferences still adheres to the standard format, with the majority, 75%, of posters using the “Introduction, methods, results, discussion” format and are heavy on text, colloquially referred to as the “wall of text” (Faulkes, 2023). The authors also demonstrated that at least 30% of these pre-selected, award winning posters had no clear reading orientation, and overall less than 50% of the posters followed good graphic design practices (Faulkes, 2023).

Given that a plethora of guidelines for poster preparation, it remains open why the quality of posters at conferences is still poor. In particular as only award-winning posters were included in the quantitative study, it stands to reason that the majority of posters have even lower overall appeal (Faulkes, 2023). Where in the poster preparation, a complex and challenging visualization format, are biological scientists struggling, do early career scientists receive training in this common visualization format?, and are they familiar with existing guidelines and helpful articles? With a qualitative survey and in-person interviews to shed light on the subjective perspective of the current struggles of life scientists in poster preparation. Using an observational, survey-based approach, we asked life scientists about their time commitment towards poster preparation, including the amount of time spend on design aspects. We also enquired on previous training they had received and the software and design process towards preparing this visual format. Our data thus summarises challenges faced by likely many more scientists and may thus help in orienting readers towards preparing a suitable training for early career scientists.

## Results

### Online User Survey

In an anonymous online survey we asked scientists working in biological and medical research 9 questions about their poster preparation practices. In total, 90 responses were collected (Figure 1A). No personal data was collected from voluntary respondents and participants were informed in writing that anonymous answers from the survey would be used to review of poster design practices, that participation was voluntary, that they could withdraw any time, and whom to reach for further questions.

**Figure 1:**
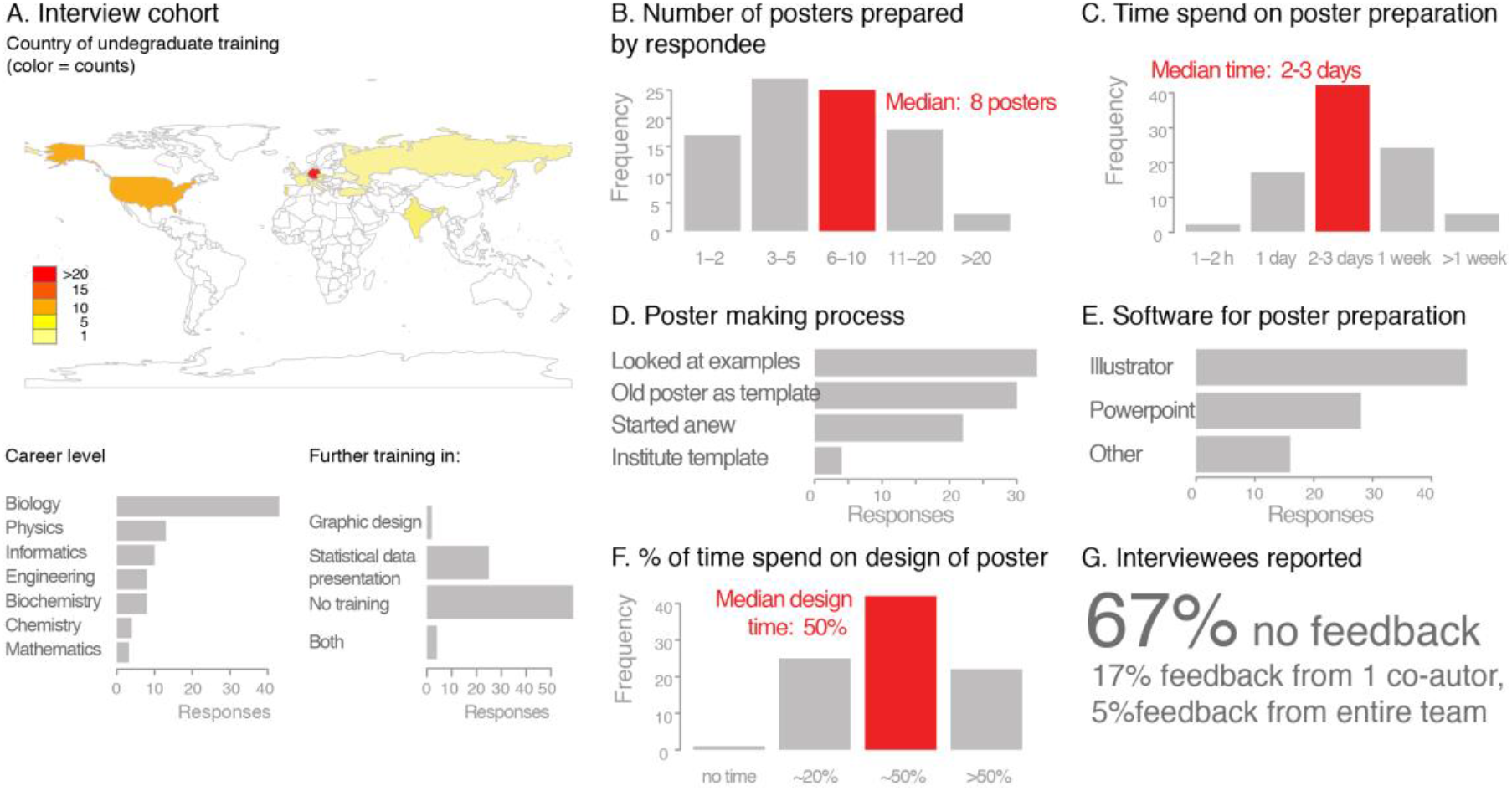
Online Survey on the poster preparation process among life scientists.

Questions of the online-survey: Country of studies, Subject of studies, Previous training in poster making, Number of posters prepared, Time spend on poster preparation, Time spend on poster design, Poster making process (designed from scratch/template), Software used for poster preparation, Feedback. Data was analysed descriptive statistics and the distribution in percentages for each response shown. Diagrams were prepared with the statistical software R and figures assembled with vector graphic software.

The online survey included responses 90 scientists who had received their undergraduate training in 26 countries (Figure 1A). Around half of the group had studied biology/related subjects, the other half included physicist, engineers, chemists and computer scientists working in biological or medical research (Figure 1A). 25 had received some training in statistical data representation, 2 respondents reported training graphical presentation of data and/or design principles for information design, and 4 had received training in both subjects (Figure 1A).

During their academic careers, respondents had already prepared a median of 8 posters (Figure 1B). They self-estimated that each poster took two to three working days in preparation time, with around half of this time, one to one-and-a-half days, exclusively devoted to design aspects (Figure 1C, F). For poster preparation, most of which were prepared without an existing template, 51% of the online polled participants used vector-based graphic software (Adobe Illustrator, CorelDraw, Inkscape) and 31% used PowerPoint (Figure 1D,E). The majority of those polled had either received no feedback or only limited feedback from just one co-author (Figure 1G).

### User Interviews

We performed in person interviews with 23 participants presenting a poster at a molecular biology conference (Figure 2A). We interviewed around 10% of the total poster presenters and were limited to the poster session times when presenters were found at their poster stands (in total two presentation slots in the course of the conference).

**Figure 2:**
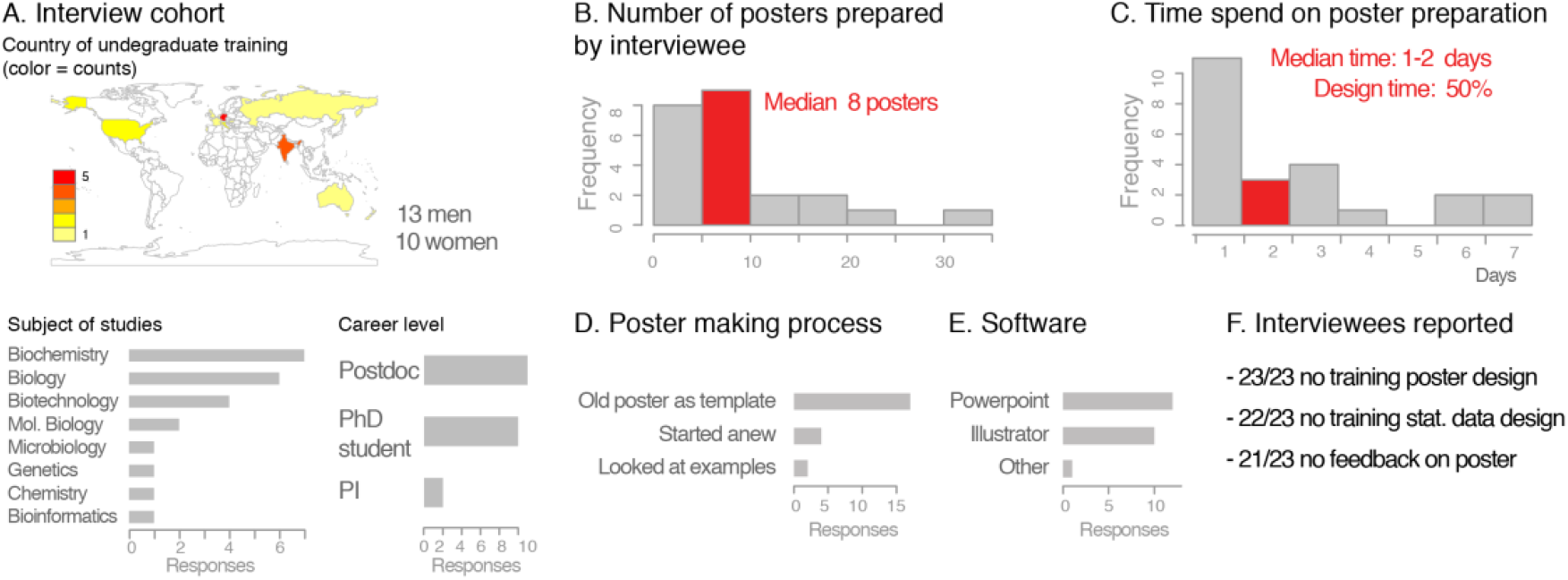
In-person interview with life scientists on their poster preparation processes.

Prior to responding to questions, interviewees were informed that anonymous answers from the survey would be used to review of poster design practices, that participation was voluntary, that they could withdraw any time, and whom to reach for further questions. No personal data was collected from voluntary respondents.

#### Questions of the in-person interview

Gender [Male/female, categorical], Career stage [Predoc/Postdoc/PI, categorical], Country of studies, Subject of studies, Time for poster preparation [<1h, 1 day, >1 day, categorical], Time spend on poster design [none, <1h, > 1h, categorical], Software used for poster preparation [free text], Number of people involved in poster preparation/their role [numerical], Poster designed from scratch/template [categorical], Previous training in poster making [free text], Personal critique of poster [free text, what went well, what was difficult/frustrating]. Data was analysed descriptive statistics and the distribution in percentages for each response shown. Diagrams were prepared with the statistical software R and figures assembled with vector graphic software.

The 23 interviewees included 13 male and 10 female scientists, the majority of which (21/23), had completed undergraduate studies in 15 different countries in biology or a closely related field in various countries (Figure 2A). Only 1 among the 23 interviewees had received training in graphical representation of statistical data. None of the respondent reported to have received training in graphic design principles or poster preparation during either undergraduate or graduate education (Figure 2F).

The career stages were diverse with two principal investigators, eleven postdocs, and ten PhD students (Figure 2A). On average the interviewees had prepared around eight posters in their scientific life, but many had prepared substantially more, up to 35 posters (Figure 2B). The median time for preparing the poster was one to two days, with half of the time spend on poster design aspects including layout, colours, and text arrangement/fonts etc (Figure 2C).

When discussing the process of poster presentation, it emerged that most interviewees (17/23) had not started their poster from scratch, but instead recycled an older poster of theirs or a colleague. In most cases this template was a PowerPoint file, which was also the software most used for poster preparation for 13 participants, while 10 used a vector-graphic software (Figure 2D).

Some interviewees, especially those who had prepared posters with PowerPoint, expressed frustration that images resolution suffered after scaling the slide to poster size (Figure 2E). Correspondingly, several participants stated that they lacked the time to learn using a vector-based software and had not been given a chance for a training in appropriate software. One person stated that the poster process took almost a week because they used the poster preparation as a chance to learn Illustrator. Similar to the online responses, the interviewees had usually not received feedback on their poster preparation from either co-workers or their principal investigator.

## Discussion

Guides and resources are published to help life scientists create powerful posters. Posters however still pose a challenging visualization format to create and are oftentimes not attractive to audiences (Faulkes, 2023). In this perspective we surveyed those preparing posters about their current challenges and approaches when designing scientific posters. This revealed that poster preparation is a laborious task, taking each scientist up to two full working days, and that one obstacle is a training deficit and the reported lack feedback.

A key observation is the large time commitment for each poster preparation. In addition, respondents report to create around eight posters as a PhD student in biology, which sums up to 14-21 working days spend on poster preparation. These posters are the basis for the first experience of early career researchers in a public scientific conversations. Despite this, poster preparation receives little explicit attention in the training of students and scholars. This lack of training and curricula may in part be due to a lack of insight into the current poster preparation process and its challenges. In addition, the rare feedback from colleagues and supervisors, seems a missed opportunity for training, especially when considering that these visualizations often are used for manuscripts.

While scientists spend considerable amount of time on posters, they and their audiences are still often not satisfied with the results. Lack of knowledge of suitable software, and lack of training in design principles, slow down this process and make it prone to errors when visual design principles (refs) are not applied or unsuitable software leads to compression artifacts. Life scientist, who in addition to posters also prepare data visualizations for manuscripts or slides, should be trained in software and design fundamentals. While several guidelines for poster preparation and presentation have been published (Barker and Phillips, 2021; Block, 1996; Brown, 1996, 1996; Erren and Bourne, 2007; Wang et al., 2022), general guidelines for preparing diagrams and figures exist from publishers (Nature Guidelines), and articles provide insights into the design process (Wong, 2011d, 2011a, 2011b), these resources rarely seem to reach the target audiences.

Integrating design into the core training of life scientists, ideally at undergraduate, but surely in graduate training, seems essential for better posters and eventually better overall visual communication of scientific results. Given that conference poster presentations are important in particular for early careers researchers to present their data for the first time in public, improving the poster quality likely will also improve the scientific discussions. And the scale is not insignificant: it is estimated that in total several million conference posters may be presented every year (Rowe, 2017), all of which could benefit from better poster design processes. Thus, designing biological data visualization and its many formats, including posters, should be an essential building block in curricula.

Re-assessing preparation for conferences is timely, given that in-person events are rapidly reemerging. Innovations established due to COVID restrictions and now the climate challenges additionally spurred innovations in conferences formats, including new forms of poster sessions (Skiles et al., 2022; Tao et al., 2021) and as equity and inclusivity of traditional conference formats are being debated (Sarabipour et al., 2021), more innovations can be expected. Posters are now often presented both in person and in virtual spaces, accompanied by poster-flash talks, or short recorded talks. The higher exposure expected from virtual poster presentations and short poster videos, often available for longer times that the conference, should be used as an opportunity to also improve the preparation of scientific posters. The BioVis community could and should help innovations in poster presentation by establishing e.g., easy to understand and implement-rules for poster design quality. Longer term, such rules could be integrated into data-driven interactive assessments of the visualizations and provide users feedback on their design quality, as it was tested for evaluating colour design practices in visualizations (InfoColorizer, see Yuan et al., 2022). A larger survey of poster design, possibly also a controlled laboratory study instead of self-reported observations, could inform exactly which quality criteria and rules are important to be included in such a tool.

As an immediate actionable measure, senior scientists can effectively improve poster quality by: 1. Providing a well-designed template as a starting point for their early career colleagues; 2. Establish a feedback routine that also discusses the visual design of the poster, and 3. Integrate free and open source vector-graphic software (e.g., Inkscape) in core training to accelerate professional design. These simple measures, and perspectively also improving training in visualization principles and tools, will improve the quality of poster visualizations at biology conference and benefit the overall science communication efficacy.

## Data Availability Statement

Data are available https://doi.org/10.5281/zenodo.7678682

## Acknowledgements

I would like to thank Lorenzo Amabili and Iris Morgenstern for reading an earlier draft manuscript. H.K.J. was supported by the MSNZ funding of the Deutsche Krebshilfe.

